# Human papillomavirus 16 E2 regulates keratinocyte gene expression relevant to cancer and the viral life cycle

**DOI:** 10.1101/461715

**Authors:** Michael R Evans, Claire D James, Molly L Bristol, Tara J Nulton, Xu Wang, Namsimar Kaur, Elizabeth A White, Brad Windle, Iain M Morgan

**Affiliations:** VCU Philips Institute for Oral Health Research, Virginia Commonwealth University School of Dentistry, Department of Oral and Craniofacial Molecular Biology, Richmond, VA 23298, USA; VCU Massey Cancer Center, Richmond, VA 23298; Department of Otorhinolaryngology, University of Pennsylvania Perelman School of Medicine, Philadelphia, PA 19104

## Abstract

Human papillomaviruses (HPV) are causative agents in ano-genital and oropharyngeal cancers. The virus must reprogram host gene expression to promote infection, and E6 and E7 contribute to this via targeting of cellular transcription factors including p53 and pRb, respectively. The HPV16 E2 protein regulates host gene expression in U2OS cells and in this study we extend these observations into TERT immortalized oral keratinocytes (NOKs) that are capable of supporting late stages of the HPV16 life cycle. We observed repression of innate immune genes by E2 that are also repressed by the intact HPV16 genome in NOKs. RNA-seq data identified 167 up and 395 downregulated genes by E2; there was a highly significant overlap of the E2 regulated genes with those regulated by the intact HPV16 genome in the same cell type. siRNA targeting of E2 reversed repression of E2 targeted genes. The ability of E2 to repress innate immune genes was confirmed in an ano-genital immortalized keratinocyte cell line, N/Tert-1. We present analysis of data from The Cancer Genome Atlas (TCGA) for HPV16 positive and negative head and neck cancers (HNC) suggesting that E2 plays a role in regulation of the host genome in cancers. Patients with HPV16 positive HNC with a loss of E2 expression exhibit a worse clinical outcome and we discuss how this could, at least partially, be related to the loss of E2 host gene regulation.

**Importance:** HPV16 positive tumors that retain expression of E2 have a better clinical outcome than those that have lost E2 expression. It has been suggested that this is due to a loss of E2 repression of E6 and E7 expression but this is not supported by data from tumors where there is not more E6 and E7 expression in the absence of E2. Here we report that E2 regulates host gene expression and place this regulation in context of the HPV16 life cycle and HPV16 positive head and neck cancers (the majority of which retain E2 expression). We propose that this E2 function may play an important part in the increased response of HPV16 positive cancers to radiation therapy. Therefore, host gene regulation by E2 may be important for promotion of the HPV16 life cycle, and also for the response of HPV16 positive tumors to radiation therapy.

## Introduction

High risk human papillomaviruses (HR-HPV) are etiological agents in ano-genital and oropharyngeal cancers (1). HPV16 is causative in around 50% of HPV positive cervical cancers (HPV+CC) and 90% of HPV positive oropharyngeal cancers (HPV+OPC), the latter of which has reached epidemic proportions over the past generation (2-5).

Following infection, HPV DNA ultimately reaches the nucleus where cellular factors induce transcription from the viral genome. The viral oncogenes E6 and E7 create an environment that promotes cellular proliferation and one aspect of this manipulation is the alteration of host gene transcription. E7 binds to pocket proteins pRb, p130 and p107 and blocks their functional interaction with E2F transcription factors resulting in transcriptional activation of their target genes (6). In addition, E7 can disrupt the DREAM complex resulting in alleviation of repression of DREAM targets and activation of B-Myb target genes (7); all of these changes induced by E7 are proliferative for the cell. The E6 protein interacts with and mediates degradation of p53 blocking the ability of this protein to regulate host gene transcription in response to cellular stress. E6 also interacts with p300 to disrupt the role of this protein in regulating p53 function and undoubtedly disrupt the function of additional p300 target proteins (8-16). Therefore, these two proteins induce a radical transcriptional reprogramming of the cell to promote the HPV life cycle.

The E2 protein of HPV16 binds to 12bp palindromic target sequences surrounding the A/T rich origin of replication and recruits the viral helicase E1 to the origin via a protein-protein interaction (17). This process initiates viral DNA replication. In addition, E2 can activate or repress transcription from the viral long control region (LCR) that incorporates the origin of replication (18). A third characterized role for E2 in the viral life cycle is to act as a viral segregation factor, binding to viral DNA and host chromatin simultaneously during mitosis to insure that the viral DNA is present in the daughter cell nuclei (19).

We have demonstrated that in U2OS cells there is a fourth function for HPV16 E2; regulation of host gene expression (20). However, the precise role of this E2 function during the viral life cycle remains ill defined. Genomic studies of high risk E2 regulation of the host genome in keratinocytes have been carried out using transient overexpression of E2 proteins using a viral delivery mechanism (21-23). While these studies have identified genes regulated by E2, it is difficult to fully delineate those genes regulated by E2, as opposed to those that are activated in response to a transient overexpression of a toxic level of E2 protein. There have been individual studies looking at the interaction between E2 and host factors, which have demonstrated that E2 can regulate host transcription via C/EBP (24), nuclear receptors (25) and AP1 (26-30). However, to date there has not been a systematic characterization of HPV16 E2 specific regulation of host gene expression in keratinocytes expressing a stable, physiologically tolerated level of E2 protein. This report provides such a characterization. In addition, we provide context for this E2 regulation in the HPV16 life cycle and also in HPV16 positive head and neck cancer (HPV16+HNC).

Previously we characterized host gene regulation by HPV16 in TERT immortalized “normal” oral keratinocytes (NOKs) (31). We developed the NOKs system for the express purpose of investigating functions in the viral life cycle that cannot immortalize keratinocytes, and E2 regulation of host gene transcription is one such function. We generated NOKs expressing E2 and investigated E2 regulation of host gene expression. In parallel we characterized the regulation of host gene transcription by the entire HPV16 genome. The results demonstrate a highly significant overlap between E2 regulated genes and those regulated by the entire HPV16 genome; the E2 regulated genes represented a sub-set of those regulated by HPV16. Many innate immune response genes were downregulated by E2 that were also downregulated by HPV16, including a signature set of genes that are ordinarily elevated for a prolonged period by the unphosphorylated interferon stimulated gene factor 3 (U-ISGF3) complex following viral exposure (32, 33). Interferon stimulation of transcription is mediated via the ISGF3 complex which is composed of phosphorylated STAT1 and STAT2 complexed with IRF9 (34). Following interferon signaling, a U-ISGF3 complex persists and is responsible for elevated expression of an innate immune gene signature set, many of them encoding known anti-viral proteins. It is proposed that this persistent elevated gene expression is a form of immune memory protecting the cell from subsequent viral infections. Repression of this gene set by HPV16 is likely an important event in promoting the viral life cycle. Others have demonstrated repression of innate immune gene expression by high risk HPV (35-38) and have assigned E6 and E7 as potential mediators of this repression (39-42). We demonstrate that E2, E6 and E7 can all contribute to repression of U-ISGF3 target genes in NOKs. siRNA targeting of E2 alleviated transcriptional repression by E2. We also demonstrate that E2 can regulate host gene transcription in ano-genital keratinocytes demonstrating that this regulation is not particular to the oral cavity. Expression of the U-ISGF3 target gene set is down regulated in HPV16 positive (HPV16+HNC) versus negative (HPV-HNC) head and neck cancers from The Cancer Genome Atlas (TCGA) (31). Repression of U-ISGF3 genes in HPV16+HNC is lost in those tumors that lacked E2 expression. Recent results demonstrate that the lack of E2 expression in HPV16+HNC results in a worse clinical outcome (43-46). We propose that host gene regulation is the fourth important function for E2 during the HPV16 life cycle (in addition to viral genome replication, transcription and segregation), and that this function is relevant to cancer outcomes.

## Results

### Generation and gene expression characterization of clonal NOK cell lines expressing HPV16 E2

Our previous studies on regulation of host gene transcription by the entire HPV16 genome utilized clonal cell lines that contained HPV16. The clonal nature of the line used for the RNA-seq analysis allowed us to identify a high number of HPV16 regulated genes whose expression changes were then validated in additional clonal cell lines. To gain a deeper understanding of HPV16 E2 regulation of host gene transcription we took the same approach and generated several clonal NOKs cell lines expressing E2 (Fig. 1a). Clone NOKs+E2-7 was used for RNA-seq analysis, and gene expression compared with NOKs only. Simultaneously, we took a NOKs+HPV16 clone and carried out RNA-seq analysis and again gene expression was compared with NOKs only. We carried out the RNA-seq with two independent samples and identified regulated genes that were significantly changed between the E2 and HPV16 containing clones and the parental NOKs in both data sets. Using a fold cut-off of 1.5 and a DESeq2 corrected p-value<0.05, E2 upregulated 167 genes and downregulated 395 genes, while the entire genome had 775 upregulated and 817 downregulated genes. Table S1 lists the regulated genes (GEO accession number GSE1216267). The known contribution of E6 and E7 to regulate host gene transcription at least partially explains the increased number of gene changes in the cells containing the entire HPV16 genome. There was a substantial overlap between the E2 regulated genes with those altered in the presence of HPV16, and this is summarized in Fig. 1b. 54 of the 167 E2 upregulated genes were also upregulated in NOKs+HPV16 (p-value = 1.2×10^35^), while 221 of the 395 downregulated genes were also downregulated in NOKs+HPV16 (p-value = 6.8×10^−210^). Table S1 includes the lists of genes that overlap between the two data sets. There was a striking number of innate immune genes targeted by both E2 and the entire HPV16 genome in NOKs. Previously we demonstrated a targeted repression of the U-ISGF3 gene set by HPV16 and this repression was retained by the E2 protein by itself (Table 1).

**Figure 1.**
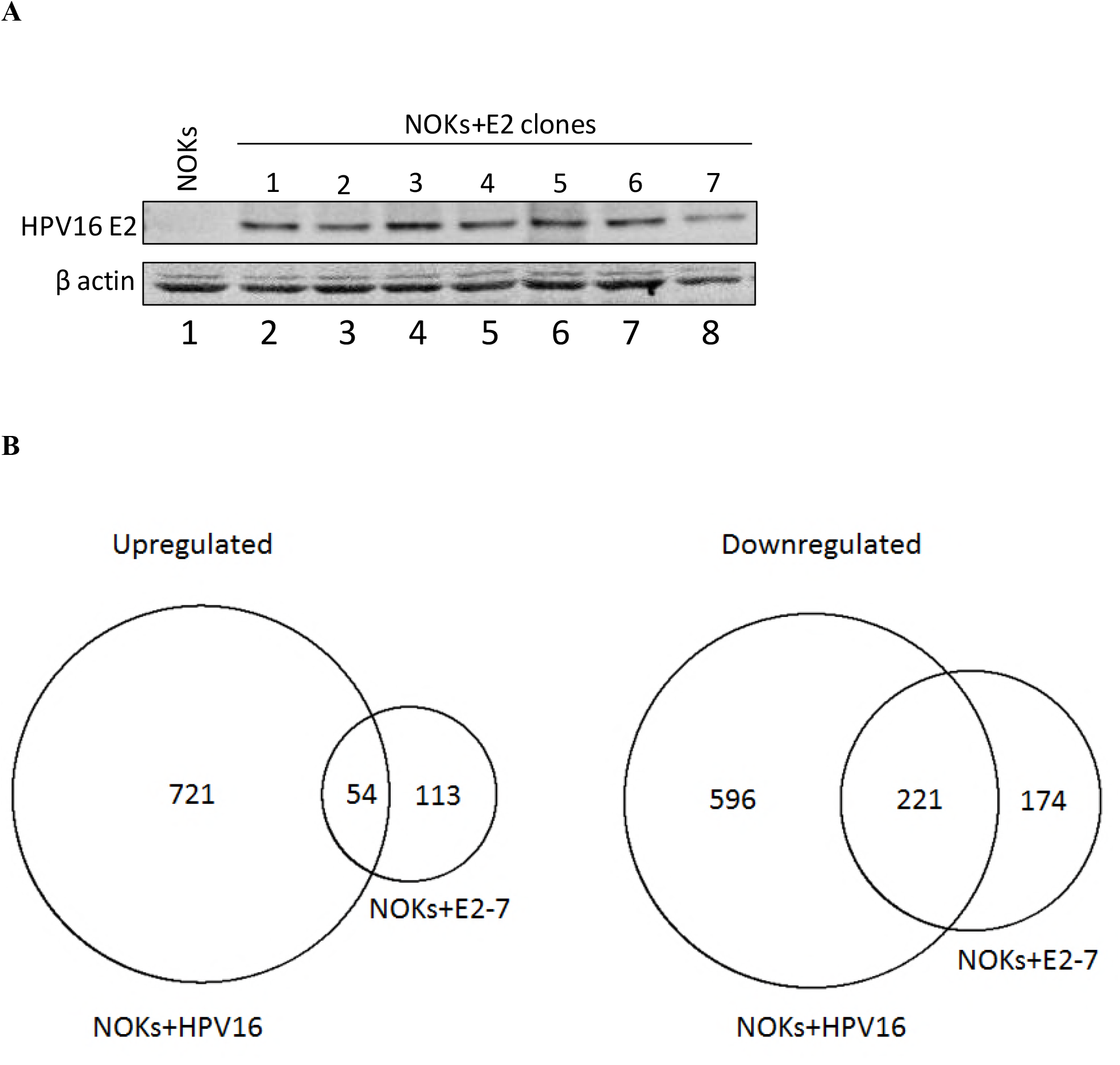
E2 regulation of host gene expression in NOKs cells. **A)** Clonal cell lines were generated by G418 selection following transfection with a HPV16 E2 expression plasmid and western blotting for E2 carried out on cell extracts. β-actin is shown as a loading control. **B)** RNA-seq was carried out with NOKs+E2-7 alongside NOKs+HPV16 (see text for details). The numbers in the circles represent the number of genes whose expression is changed 1.5 fold or greater when compared with the levels in parental NOKs. The overlap between the genes regulated by NOKs+E2-7 and NOKs+HPV16 are indicated as shared between the two data sets. The overlap of both the upregulated and downregulated genes is highly significant as described in the text. Table S1 lists the genes in each set and also those that overlap between the two samples.

**Table 1.** U-ISGF3 genes are repressed by E2 and the entire HPV16 genome in NOKs cells. If the genes were downregulated 1.5 fold or greater this is indicated with a Yes, and if not by a No.

|  | <u>NOKs+E2</u> | <u>NOKs+HPV16</u> |
| --- | --- | --- |
| **BATF2** | No | No |
| **BST2** | Yes | Yes |
| **DDX58** | Yes | Yes |
| **DDX60** | Yes | Yes |
| **EPSTI1** | No | No |
| **HERC5** | No | No |
| **HERC6** | Yes | Yes |
| **IFI27** | No | Yes |
| **IFI35** | Yes | Yes |
| **IFI44** | Yes | Yes |
| **IFI44L** | Yes | Yes |
| **IFIH1** | Yes | Yes |
| **IFIT1** | Yes | Yes |
| **IFIT3** | Yes | Yes |
| **IFITM1** | Yes | Yes |
| **IRF7** | Yes | Yes |
| **ISG15** | Yes | Yes |
| **MX1** | Yes | Yes |
| **MX2** | Yes | Yes |
| **OAS1** | Yes | Yes |
| **OAS2** | Yes | Yes |
| **OAS3** | Yes | Yes |
| **OASL** | No | No |
| **PLSCR1** | Yes | Yes |
| **RARRES3** | No | No |
| **RTP4** | No | No |
| **STAT1** | Yes | Yes |
| **TMEM140** | No | No |
| **XAF1** | Yes | Yes |

The E2 protein is over expressed in the NOKs+E2 cells compared with the levels in NOKs+HPV16, as is true in previous studies investigating host gene regulation by individual HPV proteins, such as E6 and E7. However, the highly significant overlap in the genes regulated by E2 by itself and by the entire HPV16 genome in NOKs strongly suggests that the regulation of host gene transcription by E2 is an important process in the HPV16 life cycle.

Given the conservation in U-ISGF3 gene repression between HPV16 E2 and the entire HPV16 genome, validation of the E2 RNA-seq data was focused on this gene set. The protein expression of IFIT1, MX1, STAT1 and IRF9 was determined in the NOKs+E2 clonal lines lines (Fig. 2a). In 5 of the 7 lines (1, 2, 3, 6, 7) there was a repression of MX1 and IFIT1 protein levels. The upstream activators of these genes and components of the ISGF3 complex, STAT1 and IRF9, were also downregulated in the same clones. We selected three clones for U-ISGF3 gene RNA analysis; NOKs+E2-1 and NOKs+E2-7 which had good repression of STAT1 and IRF9, and NOKs+E2-2 which had an intermediate repression of these proteins. The results are shown in Fig. 2b. All of the investigated genes were strongly repressed in NOKs+E2-1 and NOKs+E2-7 (lanes 2 and 4) while NOKs+E2-2 demonstrated an intermediate repression (lane 3). Therefore, repression correlates with downregulation of their upstream regulatory factors STAT1 and IRF9 (Fig. 2a). We next investigated expression of STAT1, IRF9 and IFNκ RNA levels in the three clones (Fig. 2c). All three repressed IFNκ expression while NOKs+E2-1 and NOKs+E2-7 had a stronger repression of STAT1 and IRF9 than in NOKs+E2-2, reflective of the protein levels of STAT1 and IRF9 in the three clones (Fig. 2a).

**Figure 2.**
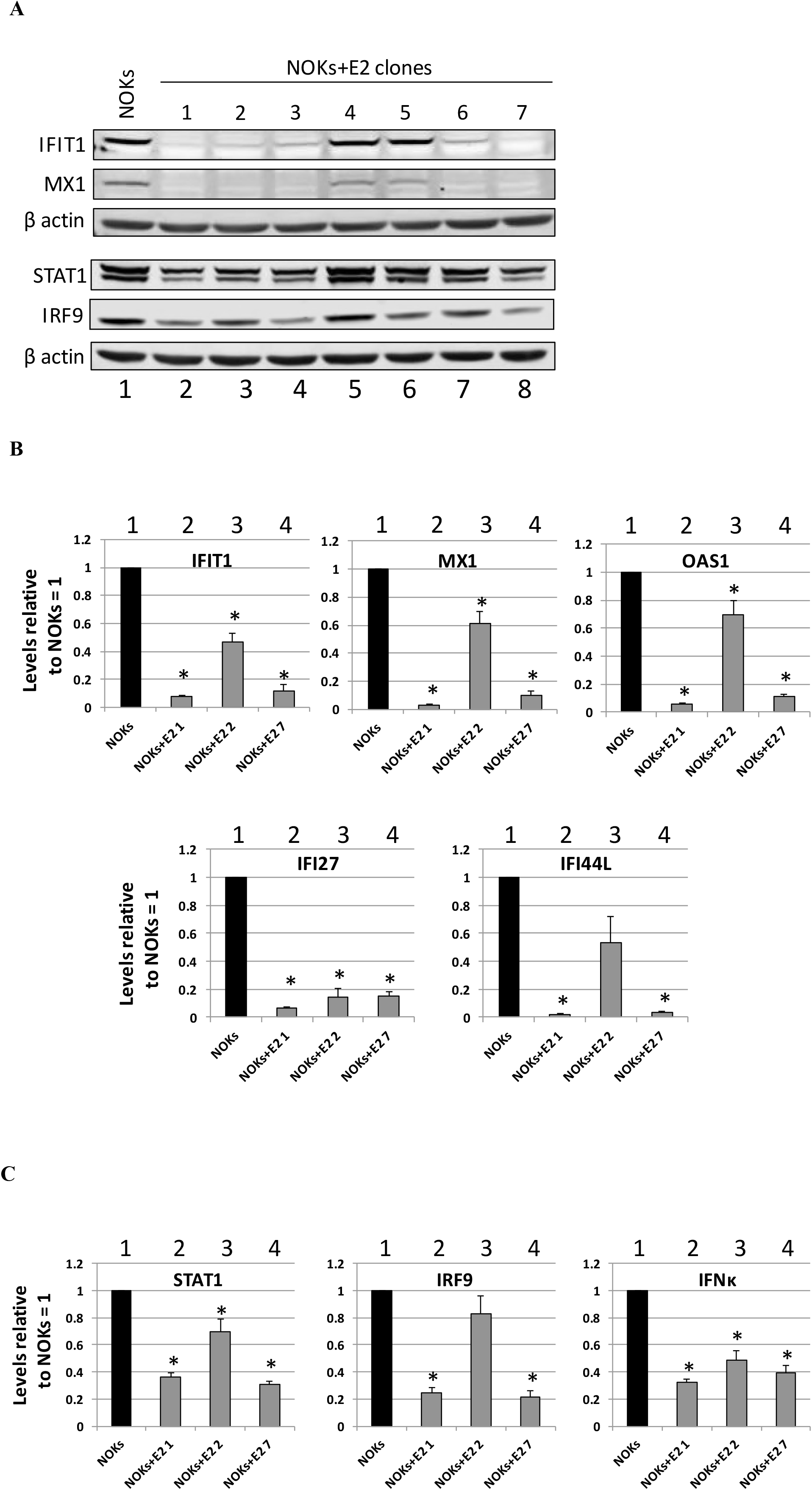
E2 represses U-IGF3 gene expression in multiple NOKs+E2 clonal cell lines. **A)** Western blots using antibodies against the indicated cellular proteins were carried out on extracts from the E2 clones. β-actin is shown as a loading control. **B)** Expression levels of a sub-set of U-ISGF3 genes in NOKs and NOKs clones expressing E2. Results are expressed as fold change from that observed in parental NOKs and represent the average of three independent experiments. **C**) Expression levels of STAT1, IRF9 and IFNκ genes in NOKs and NOKs clones expressing E2. Results are expressed as fold change from that observed in parental NOKs and represent the average of three independent experiments. Bars marked with * in B and C are significantly different from NOKs (p-value < 0.05) as determined using a student’s t-test.

Previous studies have demonstrated a role for E6 and E7 in regulation of host gene transcription, including innate immune gene repression. To investigate whether these proteins also regulated U-ISGF3 expression in NOKs cells pooled cell lines were generated expressing HPV16 E6 (NOKs+E6) and HPV16 E7 (NOKs+E7) using retrovirus transduction. In addition, we generated a pooled NOKs+E2 cell line (expressing HPV16 E2), using retroviral delivery to allow a direct comparison with the E6 and E7 expressing cells. The expression of the HA-tagged E7 and E2 proteins was confirmed by western blotting (Fig 3a, lanes 2 and 3 respectively). We could not detect the HA-tagged E6 fusion protein, presumably due to low expression levels. To confirm functional E6 expression in these cells the levels of the E6 degradation target, the p53 protein, were determined (Fig. 3b). The levels of p53 protein are down in the NOKs+E6 cells compared to control (compare lane 3 with lane 1) confirming functional E6 expression in these cells. All of the overexpressing lines expressed the appropriate viral RNA (not shown). The protein expression of two of the most repressed U-ISGF3 genes, IFIT1 and MX1, was investigated in the E2, E6 and E7 expressing NOKs (Fig. 3c). E2, E6 and E7 each downregulate expression of MX1 while E2 and E7 both repress IFIT1 but E6 does not. To confirm that this downregulation was due to gene expression levels of the IFIT1 and MX1 genes, RT-qPCR was performed on three independent RNA samples prepared from the cells and additional U-ISGF3 genes were investigated (Fig. 3d). E2 is the strongest repressor for all of the innate immune genes investigated. Reflecting the protein expression levels, E6 does not repress transcription of IFIT1. We investigated the expression of IFNκ and STAT1, a component of IFNκ’s downstream target ISGF3 complex (Fig. 3e). All three viral proteins are able to repress expression of IFNκ to a similar extent, and all three can repress expression of STAT1 although E2 is again the strongest repressor. These results demonstrate that all three viral proteins can repress U-ISGF3 target genes, potentially via targeting of the upstream activators of these genes, STAT1 and IFNκ. These results also demonstrate that the downstream targeting of U-ISGF3 genes is different between the viral proteins and do not exclusively depend upon the downregulation of IFNκ (as E6 does not repress IFNκ expression but can target downstream genes). To confirm that retroviral transduction alone did not interfere with the expression of U-ISGF3 expression, the levels of IFIT1 and MX1 proteins were determined in NOKs, NOKs+pOZHA (the vector used to generate the E2 expressing retroviral pooled cells), NOKs+pMSCV (the vector used to generate NOKs+E6 and NOKs+E7) and NOKs+pcDNA (the vector used to generate the NOKs+E2 clonal lines in Fig. 1). Figure 3f demonstrates that there is not a significant repression of either IFIT1 or MX1 with any of the vectors when compared with the expression in NOKs alone (lane 1).

**Figure 3.**
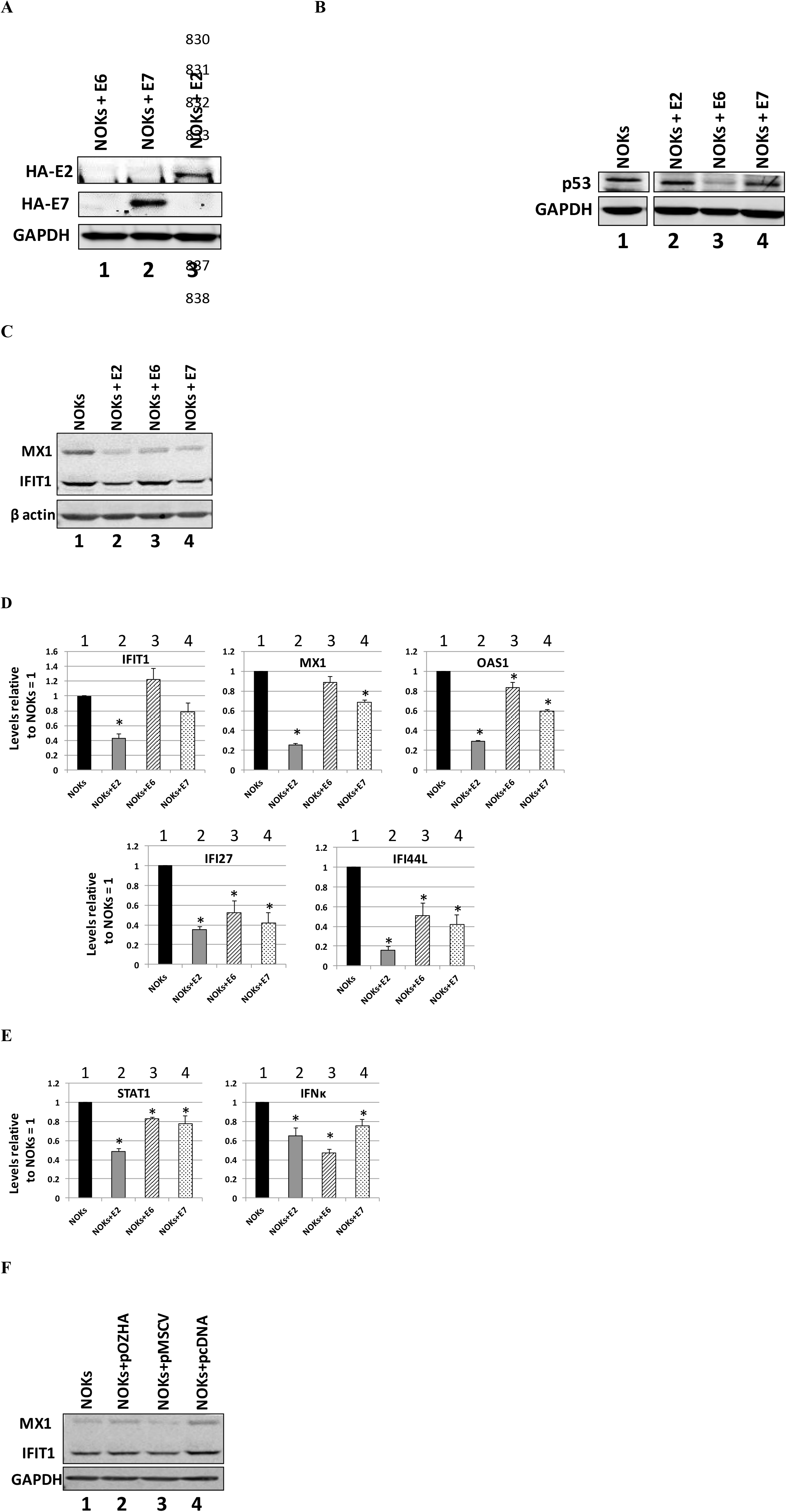
HPV16 E2, E6 and E7 can repress innate immune gene expression in NOKs. **A)** Western blot of NOKs cells transduced and selected following infection with retroviruses expressing HA-tagged E6 (lane 1), E7 (lane 2) and E2 (lane 3) with an HA antibody. GAPDH is shown as a loading control. **B)** Western blot of NOKs cells transduced and selected following infection with retroviruses expressing HA-tagged E6 (lane 1), E7 (lane 2) and E2 (lane 3) with a p53 antibody. GAPDH is shown as a loading control. There were lanes between the NOKs (lane 1) and the viral protein expressing NOKs lanes (2-4) that have been removed for clarity, but the NOKs sample image is taken from the same membrane and exposure. **C)** Western blotting of NOKs cells and NOKs expressing the indicated viral proteins with MX1 and IFIT1 antibodies. β-actin is shown as a loading control. **D)** Expression levels of a sub-set of U-ISGF3 genes in NOKs and NOKs expressing the indicated viral proteins. Results are expressed as fold change from that observed in parental NOKs and represent the average of three independent experiments. Standard error bars are shown. **E)** Expression of STAT1 and IFNκ genes in NOKs and NOKs expressing the indicated viral proteins. **F)** Transduction with retroviral vectors does not induce repression of U-ISGF3 proteins. Results are expressed as fold change from that observed in parental NOKs and represent the average of three independent experiments. Standard error bars are shown. Bars marked with * in D and E are significantly different from NOKs (p-value < 0.05) as determined using a student’s t-test.

### Transcriptional repression by HPV16 E2 is reversible

To investigate whether transcriptional repression by E2 is reversible, we used an siRNA targeting E2 (47) in NOKs+E2-1 and NOKs+E2-7 and investigated the RNA levels of E2, IFIT1, MX1 and IFNκ at the RNA level in NOKs+E2-1 and NOKs+E2-7 mock treated (1, 4, 7, 10), siRNA E2 treated (2,5,8,11) and siRNA luciferase treated (3, 6, 9, 12) (Fig. 4a). Mock and siRNA luciferase treatment resulted in identical expression levels of the genes investigated. siRNA targeting E2 reduced the expression level of E2 in both clones and resulted in an increase in the RNA expression of IFIT1, MX1 and IFNκ. Therefore, the repression by E2 is at least partially reversible. Figure 2 demonstrates that IFIT1 and MX1 are over 10 fold transcriptionally repressed by E2 but the levels of IFIT1 and MX1 following down regulation of E2 levels (Fig. 4a) increase by only 3-4 fold. This may be due to residual E2 levels present following the siRNA treatment, or that more time is required for complete relief from transcriptional repression, or that E2 induces a permanent epigenetic marker on the genes that results in constant repression irrespective of E2 levels. These possibilities will be investigated in the future. To confirm that the regulation of RNA was reflected at the protein level, western blotting for E2 and IFIT1 was carried out (Fig 4b.). As with RNA analysis, the cells were mock treated (lanes 1 and 4), treated with the E2 siRNA (lanes 2 and 5) or treated with a control siRNA targeting luciferase (lanes 3 and 6). Forty-eight hours following siRNA treatment, protein was prepared and western blotting carried out. The siRNA targeting E2 attenuated expression of this protein (middle panel) and this downregulation corresponded with an increase in IFIT1 protein levels (upper panel). This observation was reproducible. To confirm that the increased expression of IFIT1 and MX1 in the NOKs+E2 cells following downregulation of E2 expression was not due to an off target effect of the siRNA we repeated these experiments in NOKs (Fig. 4c). None of the treatments resulted in altered MX1 and IFIT1 expression levels demonstrating that there is no off target effect of the E2 siRNA that alters MX1 or IFIT1 expression.

**Figure 4.**
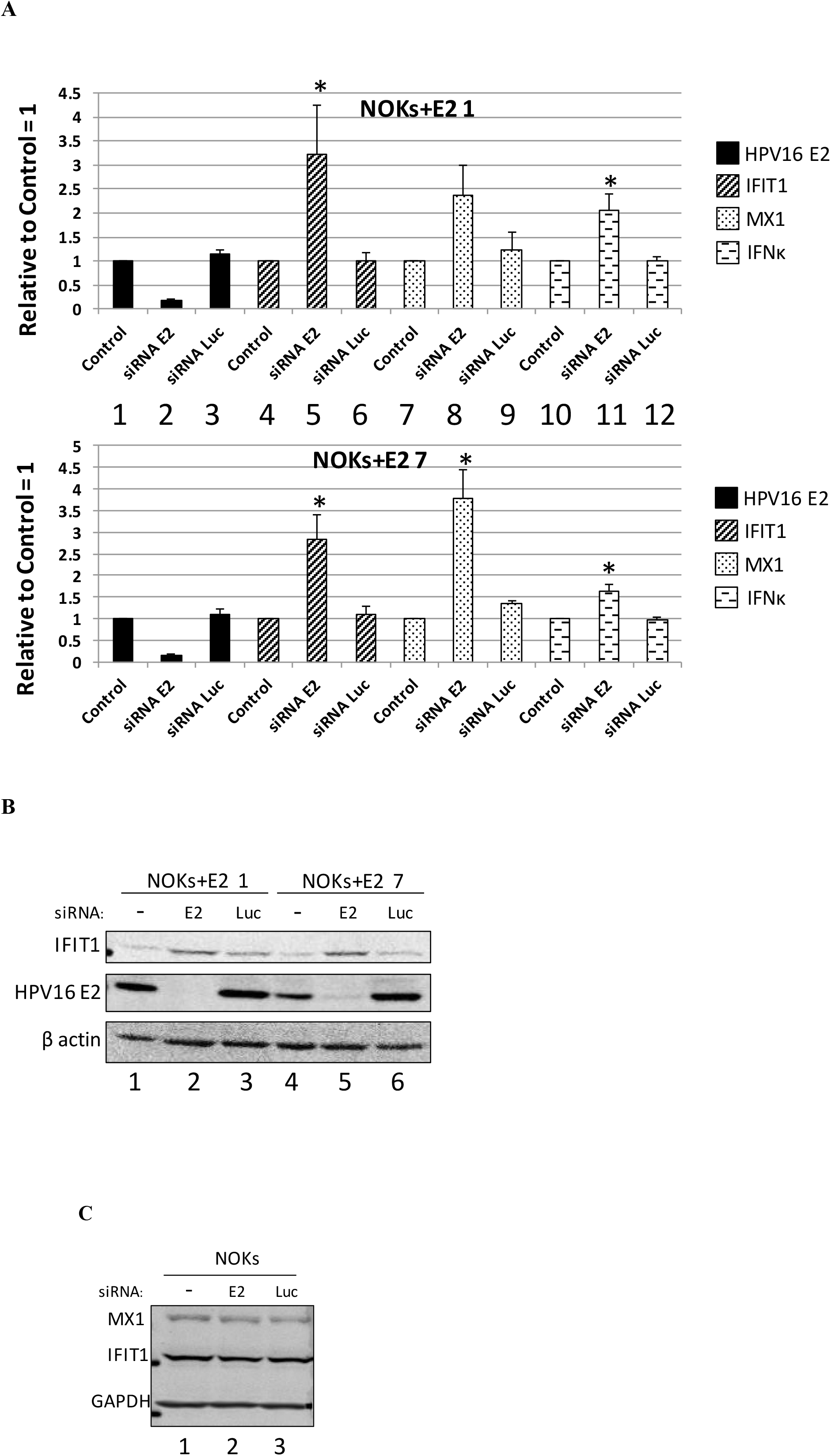
siRNA targeting of E2 partially reverses E2 repression of host genes. **A)** Expression levels of E2, IFIT1, MX1 and IFNκ genes in cells treated with the indicated siRNAs were determined. Results are expressed as fold change from that observed in each control sample and represent the average of three independent experiments. **B)**. NOKs+E2-7 and NOKs+E2-1 cells were treated with the indicated siRNAs and protein extracts prepared for western blotting. The expression levels of IFIT1 (upper panel) and E2 (middle panel) were then determined. β-actin is shown as a loading control. **C)** NOKs were mock treated (lane 1) or treated with the E2 (lane 2) or Luc (lane 3) siRNA in exactly the same way as the NOKs+E2 shown in B and protein extracts prepared for western blotting. The expression levels of MX1 (upper panel) and IFIT1 (middle panel) were then determined. GAPDH is shown as a loading control. Bars marked with * in A are significantly different from mock treated cells (p-value < 0.05) as determined using a student’s t-test.

One mechanism that HPV16 utilizes to regulate host gene transcription is to methylate host DNA (48). To investigate whether the E2 protein was regulating transcription from the host genome using this mechanism we treated cells with Decitabine (5-aza-2-deoxycytidine), a drug that reverses DNA methylation (49). The RNA levels of IFIT1 and MX1 and the upstream regulators STAT1, IRF9 and IFNκ were investigated following Decitabine treatment in NOKs, NOKs+E2-1, NOKs+E2-7 and NOKs+HPV16 (Fig. 5a). For IFIT1 Decitabine treatment in NOKs results in a small but significant increase in RNA expression (compare lane 2 with 1) while there was a much stronger response to Decitabine in the presence of E2 or the HPV16 genome. The increase in IFIT1 levels in the E2 clones (lanes 4 and 6) were statistically the same as that observed for HPV16 (lane 8). MX1 regulation is more complex; note the MX1 figure in Fig. 5a is on a log scale. Decitabine treatment resulted in a significant elevation of MX1 RNA in NOKs (compare lane 2 with 1). In both E2 clones there was over an order of magnitude increase in MX1 following Decitabine treatment (lanes 4 and 6), significantly more than that observed in NOKs. In the presence of the entire HPV16 genome there is over a two order of magnitude elevation of MX1 RNA levels (lane 8). The expression levels of IFIT1 and MX1 proteins were then investigated (Fig. 5b). In NOKs cells, treatment with Decitabine resulted in an increased expression of both MX1 and IFIT1 reflecting the RNA results (compare lane 2 with 1). Treatment of NOKs+E2-1 (lanes 3, 4), NOKs+E2-7 (lanes 5, 6) and NOKs+HPV16 (lanes 7, 8) with Decitabine resulted in an increased expression of both IFIT1 and MX1. For IFIT1, the elevated levels following Decitabine treatment in all of the cells (lane 2, 4, 6, 8) was similar suggesting that methylation by E2 and the full HPV16 genome is a major mechanism used by the virus for repressing expression of IFIT1. For MX1 the story is again different. While treatment with Decitabine increased the expression of this protein in the E2 and HPV16 expressing cells, it is not increased to levels observed in NOKs that do not express any viral protein (compare the MX1 signal in lane 2 with that in 4,6 and 8). This suggests that MX1 is controlled by methylation but additional repression mechanisms in cells expressing E2 and HPV16 are involved. Therefore, the results in Fig. 5a & 5b suggest that methylation is not the only mechanism that E2 or the entire HPV16 genome uses to repress transcription of MX1. The elevated response to Decitabine for the entire genome versus the E2 clones following Decitabine treatment also demonstrates that HPV16 has, in addition to E2, additional mechanisms for repressing MX1 transcription. A role for E6 and E7 in this repression seems likely. STAT1, IRF9 and IFNκ RNA levels are all elevated following Decitabine treatment in NOKs, both E2 expressing clones and that containing the entire HPV16 genome (Fig. 5a). For both STAT1 and IRF9 the increase in expression following Decitabine treatment is higher in the E2 clones and with the HPV16 clone and the increase is similar with both E2 and HPV16. However, the fold increase in IFNκ expression following Decitabine treatment is no different between NOKs and the E2 clones and cells containing HPV16. This suggests that there is not a direct role for E2 induced methylation in repressing these genes in the NOKs.

**Figure 5.**
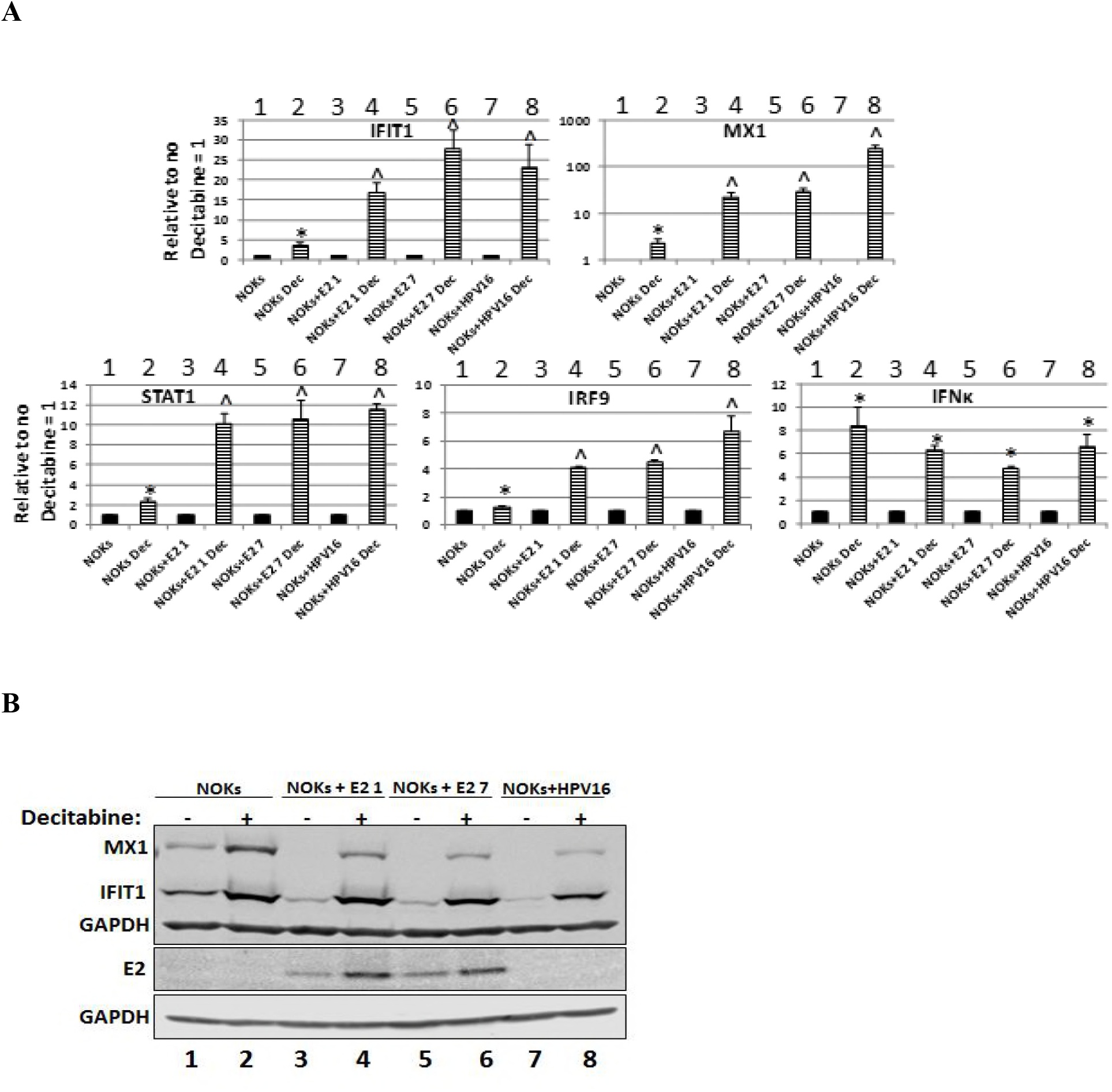
Transcriptional repression by E2 and the entire HPV16 genome is relieved by treatment with the DNA de-methylase Decitabine. **A)** Expression levels of several innate immune response genes following Decitabine treatment of the indicated cells. Results are expressed as fold change over the no Decitabine treatment sample in each case and represent the average of three independent experiments. In A, bars marked with * are significantly elevated from levels in Decitabine non-treated cells. Those marked ^ are significantly elevated from levels in Decitabine non-treated cells and the fold change in elevation is significantly greater than that in NOKs. A student’s t-test was used. **B)** Western blots for IFIT1 and MX1 on protein extracts from the indicated cells with and without Decitabine treatment as indicated. GAPDH is shown as a loading control.

To confirm that the alleviation of repression with Decitabine was not due to a reduction in E2 protein levels, western blotting was carried out to determine the E2 protein levels following Decitabine treatment (Figure 5b, lower panels). Decitabine actually increases the levels of the E2 protein, perhaps indicating that the promoter driving E2 expression is also methylated in the NOKs+E2 cells.

### HPV16 E2 can also repress host gene transcription in ano-genital keratinocytes

To investigate whether the E2 protein could also regulate transcription from the host genome in ano-genital keratinocytes, HPV16 E2 was over expressed in N/Tert-1 cells. N/Tert-1 were generated by immortalizing foreskin keratinocytes with the telomerase enzyme (50). A retroviral expression vector encoding E2 was used to generate N/Tert-1+E2. Fig. 6a demonstrates expression of the E2 protein in N/Tert-1 cells (compare lane 2 with 1), and also shows that both IFIT1 and MX1 protein expression levels are acutely repressed by E2 in this cell line (compare lane 4 with 3 and 6 with 5). We next investigated expression of a sub-set of U-ISGF3 genes in the N/Tert-1+E2 versus control cells (Fig. 6b) and 4 out of 5 genes were repressed by the E2 expression. Although IFI27 was greatly repressed by E2 in NOKs cells (Figs. 1e and 2c), it was not repressed by E2 in the N/Tert-1. Finally, we investigated whether E2 could also repress expression of components of the ISGF3 complex and its upstream activator IFNκ (Fig. 6c). STAT1 expression is repressed by E2 in these cells but the repression of IFNκ expression is small compared with that observed in NOKs (Fig. 2d). The IFI27 and IFNκ results demonstrate that the pattern of repression of innate immune genes by E2 is not universal across all keratinocyte cell types.

**Figure 6.**
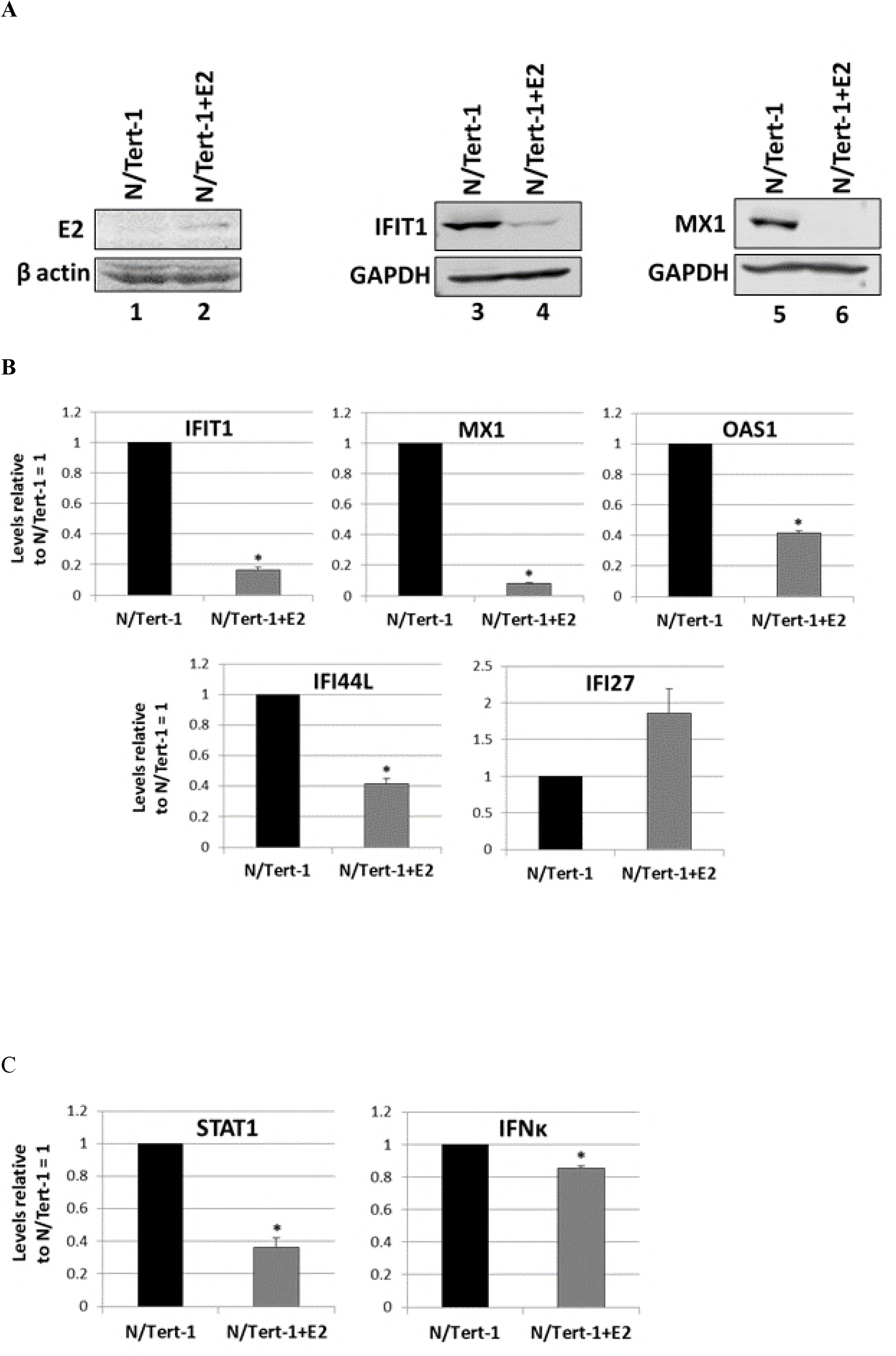
E2 also targets innate immune gene repression in NTERT, a TERT immortalized foreskin cell line. **A)** Expression of HA tagged E2 in the NTERT detected by western blotting of protein extracts with an E2 antibody. β-actin is shown as a loading control. Western blots for IFIT1 and MX1 on protein extracts from the E2 positive and negative cells. GAPDH is shown as a loading control. **B)** Expression of a sub-set of the U-ISGF3 gene set in the NTERT and NTERT+E2 cells. Results are expressed relative to NTERT levels equaling 1 and are representative of three independent experiments. **C)** Expression levels of STAT1 and IFNk genes in NTERT and NTERT+E2. Results are expressed relative to NTERT levels equaling 1 and are representative of three independent experiments. Bars marked with * in B and C are significantly different from NOKs (p-value < 0.05) as determined using a student t-test.

### A role for E2 in regulating innate immune gene expression in HPV16 positive head and neck cancers (HPV16+HNC)

Given the significant overlap between the E2 regulated genes with that of the entire HPV16 genome (Fig. 1b), we investigated whether the E2 regulated genes (Table S1) were also regulated in HPV16 positive versus negative head and neck cancers. In order to do this, we divided the head and neck cancer tumor gene expression data from The Cancer Genome Atlas (TCGA) into HPV16 positive and HPV16 negative (HPV16+HNC versus HPV-HNC). The gene changes between the HPV16 positive and negative cancers were then determined and Table 2 lists the number of gene changes identified. For the E2 upregulated genes, 29 out of 167 were upregulated by HPV16 in the tumor data set, and 137 of 395 E2 downregulated genes were downregulated by HPV16; both of these numbers are statistically significant. Next, we divided the HPV16+HNC into those that expressed E2 RNA and those that did not, {HPV16(E2+)+HNC and HPV16(E2-)+HNC}, and compared gene changes versus HPV-HNC tumors. Again, Table 2 demonstrates that there is a significant overlap between the E2 regulated genes in NOKs and the genes regulated by HPV16(E2+)+HNC and also HPV16(E2-)+HNC when compared with HPV-HNC. In the absence of E2 there is a reduction in the number of HPV16+HNC regulated genes when compared with HPV-HNC; this may be due to the reduction of the number of tumors used to generate the data (of the 60 HPV16+HNC, 16 had no E2 expression). Therefore, we cannot conclude that there is a loss of E2 gene regulation following loss of E2 expression in these tumors. However, it was notable that the repression of the U-ISGF3 gene set was lost following the loss of E2 expression (Table 3). These results strongly suggest that the host gene changes induced by HPV16 E2 to promote the viral life cycle are physicologically relevant based on similar effects manifested in HPV16+HNC. Table S2 lists the genes summarized by numbers in Table 2.

**Table 2.** The intersection between E2 genes regulated in NOKs and those regulated by the entire HPV16 genome in NOKs, and in HPV16 positive head and neck cancers. There is a highly significant overlap in the number of genes regulated by E2 and the entire HPV16 genome in NOKs. In addition, there is also a significant overlap between the E2 regulated genes and those regulated by HPV16 in head and neck cancer (HNC). The latter gene sets were determined by comparing the HPV16 positive cancers with HPV negative. The cancer data was based on analysis of The Cancer Genome Atlas head and neck cancer samples. HPV16(all)+HNC v HPV16-HNC = all of the data from HPV16 positive HNC versus that from HPV negative. HPV16(E2+)+HNC v HPV16-HNC = all of the data from HPV16 positive HNC that retain E2 RNA expression versus that from HPV negative. HPV16(E2-)+HNC v HPV16-HNC = all of the data from HPV16 positive HNC that have no E2 RNA expression versus that from HPV negative. Up = the number of genes in the data set increased 1.5 fold or higher. Down = the number of genes in the data set decreased 1.5 fold or higher. The overlap in the gene sets is particularly significant in the repressed gene set.

|  | Up | Intersection<br>with<br>NOKs+E2-7 | p-value | Down | Intersection<br>with<br>NOKs+E2-7 | p-value |
| --- | --- | --- | --- | --- | --- | --- |
| NOKs+E2-7 | 167 |  |  | 395 |  |  |
| NOKs+HPV16 | 775 | 54 | $1.2 \times 10^{-35}$ | 817 | 221 | $6.8 \times 10^{-210}$ |
| HPV16(all)+HNC<br>v HPV16-HNC | 2361 | 29 | 0.015 | 2713 | 137 | $2.4 \times 10^{-28}$ |
| HPV16(E2+)+HNC<br>v HPV16-HNC | 2427 | 30 | 0.012 | 2625 | 132 | $5.5 \times 10^{-27}$ |
| HPV16(E2-)+HNC<br>v HPV16-HNC | 649 | 12 | 0.004 | 1304 | 64 | $2.3 \times 10^{-9}$ |

**Table 3.** Repression of U-ISGF3 gene expression is lost in E2 negative HPV16+HNC. If the genes were downregulated 1.5 fold or greater this is indicated with a Yes, and if not by a No.

|  | <u>HPV16+E2-HNC</u> | <u>HPV16+E2+HNC</u> |
| --- | --- | --- |
| BATF2 | No | No |
| BST2 | No | No |
| DDX58 | No | Yes |
| DDX60 | No | Yes |
| EPSTI1 | No | No |
| HERC5 | No | No |
| HERC6 | No | Yes |
| IFI27 | No | Yes |
| IFI35 | No | No |
| IFI44 | No | Yes |
| IFI44L | No | No |
| IFIH1 | No | No |
| IFIT1 | No | Yes |
| IFIT3 | No | Yes |
| IFITM1 | No | No |
| IRF7 | No | No |
| ISG15 | No | Yes |
| MX1 | No | Yes |
| MX2 | No | Yes |
| OAS1 | No | No |
| OAS2 | No | Yes |
| OAS3 | No | No |
| OASL | No | Yes |
| PLSCR1 | No | No |
| RARRES3 | No | No |
| RTP4 | No | No |
| STAT1 | No | No |
| TMEM140 | No | No |
| XAF1 | No | Yes |

## Discussion

We have reported that NOKs can support late stages of the HPV viral life cycle as demonstrated by E1^E4 and E2 protein expression in the differentiated layers of organotypically rafted cells, along with amplification of the HPV16 genome in the same cellular layer as measured by fluorescence in situ DNA hybridization (31). We characterized the host gene regulation by HPV16 in this model system. Among the repressed gene set was a 29 innate immune gene set activated by U-ISGF3, a complex consisting of STAT1, STAT2 and IRF9 that activates transcription of these genes irrespective of interferon stimulation (32, 33). The U-ISGF3 gene set consist of a number of known anti-viral genes, therefore repression of this gene set by HPV16 is likely required for the viral life cycle. The purpose of developing and characterizing this HPV16 life cycle model in an immortalized cell line was to allow investigation of events in the life cycle that would not immortalize primary keratinocytes, such as expression of the E2 protein by itself. We generated NOKs cells expressing E2, E6 and E7 proteins and focused on the expression of U-ISGF3 genes. All three viral proteins can repress the upstream activator of ISGF3, IFNκ, while E2 is the strongest repressor of STAT1 (Fig. 3). E2 is also the strongest repressor of the U-ISGF3 target genes investigated, and E6 can only repress a sub-set of these genes. These results demonstrate that there are three viral proteins that can repress innate immune genes; E2, E6 and E7. The repression of these genes by E6 and E7 has been characterized previously (39-42), this is the first time such an activity has been credited to E2. It is possible that the three viral proteins can synergistically repress innate immune genes via different mechanisms. It is also possible that, at different points in the viral life cycle, the role of the viral proteins in repression of these genes in important. For example, following infection it has been suggested that during establishment E2 expression is strong compared with E6 and E7 and therefore repression by E2 may be important during establishment of an HPV16 infection (51).

To characterize host gene reprograming by E2 in NOKs, we generated clonal cell lines expressing this protein to be used in RNA-seq studies (Fig. 1). Using this approach with the entire HPV16 genome, an in depth understanding of host gene regulation by HPV16 was obtained and an in depth understanding of E2 host regulation was also obtained (Table S1).In 5 of the 7 E2 clones generated there was repression of ISGF3 components and U-ISGF3 targets at both the transcriptional and protein levels (Fig. 2). The reason for the failure of the two E2 clones that do not regulate transcription of these genes could be due to transfection of the cells at different points of the cell cycle, or it may be related to actual differences in the transfected cells that do not allow E2 programing of innate immune genes. To our knowledge clonal analysis of E6 and E7 over expression has not been reported, therefore it is possible that this resistance to reprograming may not be unique to the E2 protein. However, there is no doubt that E2 regulates transcription of these host genes given the repression observed in pools of E2 expressing cells from different sources (Figs. 3 and 6), and the ability to reverse this repression (Figs. 4 and 5). RNA-seq analysis demonstrated a highly significant overlap between E2 regulated genes and those regulated by the entire HPV16 genome in NOKs emphasizing the potential importance of E2 host gene regulation during the viral life cycle. MMP9 is activated by cottontail rabbit papillomavirus (CRPV) E2 protein (28), but in our data set we observed that this gene was repressed by HPV16 E2. This could be due to a difference between the HPV16 and CRPV E2 proteins or may be related to the model systems used. The U-ISGF3 complex is also targeted for repression by HPV16 E2 in the anogenital cell line N/Tert-1 (Fig. 6) demonstrating that this repression is not something that is particular to NOKs or oral keratinocytes. The repression by E2 is not identical between the two epithelial cell lines. IFNκ is poorly repressed by E2 in N/Tert-1 but well repressed in NOKs. The U-ISGF3 target gene IFI27 is well repressed by E2 in NOKs but is not repressed by E2 in N/Tert-1; if anything E2 elevates IFI27 expression in these cells. These results suggest that E2 can regulate host gene transcription differently in different anatomical site keratinocytes and this could have an important role in the viral life cycle in these different tissues.

The mechanisms that E6 and E7 use for regulation of host gene transcription are at least partially understood. The E2 mechanisms contributing to host gene transcription regulation are ill defined. Although there are target sequences for E2 in our genome, and E2 can bind to these sequences, there is no evidence that direct binding by E2 to those sequences plays a substantial role in E2 regulation of host gene transcription (52, 53). We have shown that, in U2OS cells, failure to bind Brd4 results in a loss of some E2 upregulated genes (54). This is presumably due to a protein-protein interaction recruiting E2 to Brd4 target genes. Here we demonstrate that E2 repression of transcription is at least partially reversible (Fig. 4). However, the elevation of IFIT1 and MX1 transcription following elimination of E2 expression via siRNA targeting is only 2-3 fold (Fig. 4b) whereas the repression of these genes is 10 fold and over (Fig. 2c). This could be due to the timing of the siRNA experiments, it may be that long term elimination of E2 expression would result in the return of wild type IFIT1 and MX1 expression. However, the results with Decitabine (Fig. 5) suggests that E2 promotes an active methylation of the repressed innate immune gene promoters as repression is alleviated in the presence of this drug. The repression relief is also observed in NOKs but not to the same extent as with E2. In addition to methylation, E2 also will have other mechanisms for repressing transcription but the Decitabine results do suggest a partial role for enhanced methylation. It may not be a direct effect on the promoters of the genes under study, it could be related to a transcription factor in common to the promoters under study that is regulated by E2 methylation of its gene. The Decitabine experiment also reveal a differential regulation of U-ISGF3 target genes by E2 and the entire HPV16 genome. IFIT1 levels increase equivalently in NOKs+E2-1, NOKs+E2-7 and NOKs+HPV16 following Decitabine treatment suggesting that E2 is a major repressor of these genes during the viral life cycle. However, the pattern with MX1 is different. In both E2 clones there is an order of magnitude increase in MX1 levels following Decitabine treatment, but in NOKs+HPV16 this is two orders of magnitude. Therefore, there is differential regulation of the U-ISGF3 genes by the virus and the results with MX1 suggest the virus uses additional mechanisms to repress MX1 repression outside of E2 expression. A role for E6 and E7 to synergize in this repression seems likely. MX1 is the most tightly repressed U-ISGF3 gene by HPV16. While it is known that IFIT1 can repress HPV18 replication (55, 56) (and also plays a role in HPV16 replication, Evans et al. in preparation) the role of MX1 in the viral life cycle is not known but given the level of repression, it is worthy of investigation.

Expression of E2 predicts a better survival rate in HPV16+HNC, presumably due to an increased response to radiation therapy in the presence of E2 (43-46). It is striking that loss of E2 expression in HPV16+HNC has both reduced survival and also a loss of U-ISGF3 repression when compared with HPV negative HNC. A subset of U-ISGF3 genes belongs to the interferon related DNA damage resistance signature (IRDS) that was first identified as an upregulated gene set in a head and neck cancer cell line resistant to radiation treatment (57). Elevated expression of the IRDS gene set in breast cancer predicts worse clinical outcome (58). The E2 protein can downregulate expression of IRDS genes in NOKs and this repression is exhibited in the E2 positive HPV16+HNC. Therefore, the loss of E2 expression results in elevation of a gene set, IRDS, whose increased expression predicts a worse clinical outcome in cancer. The precise mechanism of how IRDS genes regulate the response to cancer therapy is not clear, however we propose that loss of repression of this gene set following loss of E2 expression is at least partially responsible for the worse clinical outcome in HPV16+HNC that have no E2 expression.

In conclusion, we have demonstrated that E2 can regulate transcription from the host genome in both NOKs and N/Tert-1 although the regulation is different between the two keratinocyte cells. We also provide evidence that this regulation by E2 is important in the context of the HPV16 life cycle as there is a significant overlap between the E2 regulated genes and those regulated by the entire HPV16 genome in NOKs. The entire HPV16 genome regulates many more host genes due to the contribution of additional mechanisms mediated by other viral proteins such as p53 and pRb. Finally, we propose that the host gene regulation by E2 in keratinocytes has direct relevance in HPV16 cancers as loss of E2 expression promotes a worse clinical outcome combined with a significant loss of proposed E2 regulated genes in HPV16+HNC. Future work will focus on establishing the mechanisms that E2 uses to regulate host gene transcription to enhance our understanding of the viral life cycle and also the therapeutic response of HPV16+HNC and other HPV16 tumors to radiation therapy. These mechanisms could include regulation of gene splicing, regulation of miRNA expression targeting certain gene sets as well as regulation of long non-coding RNAs (lncRNAs).

## Materials and Methods

### Cell lines

Normal oral keratinocytes (NOKs) were immortalized with telomerase reverse transcriptase (TERT) (59). NOKs cells stably overexpressing HPV16 proteins (NOKs+E2, NOKs+E6, NOKs+E7) were generated by infecting NOKs with lentivirus expressing HPV16 E2 (pOZ-N-HA16E2), HPV16 E6 (p6661 MSCV-IP N-HA only 16E6 – Addgene plasmid # 42603), or HPV16 E7 (p6640 MSCV-P C-FlagHA 16E7-Kozak - Addgene plasmid #35018). Lentivirus was produced in 293T cells using packaging vectors psPAX2 and pMD2.G. NOKs+E2 clones were generated by plating 1×10^6^ NOKs on 10cm plates and transfecting with 1μg E2 expression vector (18) using Lipofectamine 2000 (Invitrogen). Transfected cells were selected with 150μg/mL G418 (Corning) with media changed every 3 days. Individual colonies surviving selection were harvested by trypsinization using cloning rings and expanded into multiple NOKs+E2 clonal cell lines. E2 expression was confirmed by qRT-PCR and western blotting. Generation of NOKs+HPV16 clones was described previously (31). N/Tert-1+E2 were prepared using the plasmids described in (60) to generate retroviruses for infection of N/Tert-1 as described (61).

### Cell Culture

Cells were grown in keratinocyte serum-free medium (K-SFM) (Gibco) with 1% (v/v) penicillin/streptomycin mixture (Life Technologies) containing 4μg/mL hygromycin B (Sigma) at 37°C in a 5% CO_2_/95% air atmosphere and passaged every 3-4 days. Cells were routinely checked for mycoplasma contamination. NOKs+E2 clones and NOKs+HPV16 clones were grown in medium containing 150μg/mL G418 (Corning). NOKs+E2, NOKs+E6, and NOKs+E7 and NTERT+E2 pools were grown in medium containing 5 μg/mL puromycin (Corning).

### Western Blotting

Cells were trypsinized, washed with PBS, pelleted, and then resuspended in 50μl of lysis buffer (0.5% Nonidet P-40, 50mM Tris, ph 7.8, 150mM NaCl) supplemented with protease inhibitor (Roche Molecular Biochemicals) and phosphatase inhibitor cocktail (Sigma). The cell and lysis buffer mixture was incubated on ice for 30 min, centrifuged for 20 min at 184,000rfc at 4°C, and supernatant was collected. Protein levels were determined utilizing the Bio-rad protein estimation assay. Equal amounts of protein were boiled in in 2x Laemmli sample buffer (Bio-rad). Samples were then loaded into a Novex 4-12% gradient Tris-glycine gel (Invitrogen), run at 100V for approximately 2 hours, and then transferred onto nitrocellulose membranes (Bio-rad) at 30V overnight using the wet blot method. Membranes were blocked in Odyssey blocking buffer (diluted 1:1 with PBS) at room temperature for 6 h and probed with relevant antibody diluted in Odyssey blocking buffer overnight at 4°C. Membranes were then washed with PBS supplemented with 0.1% Tween (PBS-Tween) before probing with corresponding Odyssey secondary antibody (goat anti-mouse IRdye800cw or goat anti-rabbit IRdye680cw) diluted 1:10,000 for 1h at 4°C. Membranes underwent washing in PBS-Tween before infrared scanning using the Odyssey CLx Li-Cor imaging system. The following antibodies were used for western blot analysis at 1:1000 dilutions in Odyssey blocking buffer (diluted 1:1 with PBS): IFIT1, MX1 (D3W7I), IRF9 (D8G7H) from Cell Signaling Technology. STAT1 (sc-346), β-actin mouse (sc-81178), β-actin rabbit (sc-130656), GAPDH (sc-47724) from Santa Cruz Biotechnology. HPV16 E2 (TVG 261) from Abcam.

### SYBR Green qRT-PCR

Cells were harvested by trypsinization and washed twice with PBS. RNA was immediately isolated using the SV Total RNA Isolation System (Promega) following the manufacturer’s instructions. Two micrograms of RNA were reverse transcribed into cDNA using the High Capacity Reverse Transcription Kit (Applied Biosystems). cDNA and relevant primers were added to PerfeCTa SYBR Green FastMix (Quantabio) and real-time PCR performed using 7500 Fast Real-Time PCR System (Applied Biosystems). Results shown are the average of three or four independent experiments with relative quantity of genes determined by the ΔΔCt method normalized to the endogenous control gene GAPDH. Primers: GAPDH, IFIT1, MX1, OAS1, IFI27, IFI44L primers used were designed by Qiagen (QuantiTech primer assay). Other primer pairs used in this study are as follows: STAT1 (Invitrogen): 5’- CAGCTTGACTCAAAATTCCTGGA-3’ (forward) and 5’- TGAAGATTACGCTTGCTTTTCCT-3’ (reverse). IRF9 (Invitrogen): 5’- GCCCTACAAGGTGTATCAGTTG-3’ (forward) and 5’-TGCTGTCGCTTTGATGGTACT-3’ (reverse). IFN-k (Invitrogen): 5’-GTGGCTTGAGATCCTTATGGGT-3’ (forward) and 5’- CAGATTTTGCCAGGTGACTCTT-3’ (reverse). E2 (Invitrogen) 5’- TGGAAGTGCAGTTTGATGGA-3’ (forward) and 5’-CCGCATGAACTTCCCATACT-3’ (reverse).

### RNA-seq analysis

RNA sequencing was carried out by the Genomics, Epigenomics and Sequencing Core at the University of Cincinnati. The methods used for RNA production and analysis to generate the lists of differentially expressed genes have been described previously (31).

### siRNA Treatment

1.5×10^5^ cells were plated onto 6 well plates. The next day cells were transfected with 25nm siRNA targeting E2 (GCAGUUUGAUGGAGACAUAUU-Dharmacon) (15) or control siRNA targeting luciferase (Ambion) using Lipofectamine 2000 (Invitrogen). 16 hours later, transfected cells were washed with PBS and media replaced. 48 hours post-wash cells were harvested by trypsinization and processed for qRT-PCR or western blotting as described above.

### Decitabine Treatment

2.5×10^5^ cells were plated onto 10cm plates. The following day 1 μM Decitabine (Abcam - ab120842) was added to media of cells. 96 hours later, cells were harvested by trypsinization and processed for qRT-PCR or western blotting as described above.

### Differential Expression in NOKs

Methods used to analyze RNA-seq data for NOK, NOK HPV16 and NOK HPV16 E2 are described in (31). The latest human assembly, Homo_sapiens.GRCH38.dna_sm.primary_assembly.fa from Ensembl along with its corresponding .gtf file was used to create a reference file using bowtie2 build command. RNA-seq fastq files for NOKs, NOKs+HPV16, and NOKs+HPV16 E2 were locally aligned to GRCh38 reference sequence using bowtie2, mapped reads were sorted by Samtools and the resulting BAM files were analyzed by DESeq2. Gene-level analysis was performed and log2 fold change was converted to ratio of NOKs HPV+/NOKs HPV-. Genes were considered differentially expressed if the adjusted p-value computed by DESeq2 was < 0.05 and the expression differences were > 1.5 fold in magnitude.

### Differential Expression in TCGA

RNA-seq Version 2 head and neck squamous cell carcinoma (HNSC) expression data was obtained from The Cancer Genome Atlas (TCGA) as described previously (46). 508 patients were used for the analysis; all were either HPV16 negative (n=448) or HPV16 positive (n =60). The RSEM expected counts for each patient/gene were rounded to the nearest whole number and matrix was analyzed by DESeq2. Gene level analysis with default parameters was performed with the same method as stated above. A Benjamini and Hochberg correction with a fixed FDR of 0.001 was used and genes with expression differences >1.5 fold in magnitude were considered differentially expressed.

### Determination of E2 status for TCGA cohort

Unmapped human reads for 60 HPV16 positive head and neck patients from TCGA were used to define raw count levels of HPV (46). Read counts were normalized by total reads/million. Levels of E2 protein were defined using the HPV16 genome coordinates (K02718.1) 3625-3800bp as this region has no overlap between E2/E4.

Tumors were assigned E2 (n = 44) or no E2 (n = 16) status by plotting E2 levels and determining a cut-off value based on change in slope. This method defined the no E2 group as no or very little expression of E2 and E2 levels are presented in supplementary data.

### P-value calculations for comparing gene expression changes

P-values for intersecting results in the bioinformatic data listed in Table 2 were obtained using the hypergeometric distribution. Calculations were based on a total of 20532 genes.

## Acknowledgements

This work was supported by VCU Philips Institute for Oral Health Research and the National Cancer Institute Designated Massey Cancer Center grant P30 CA016059 and by the grant R03 DE026230 from the National Institute for Dental and Craniofacial Research.

## References

1. zur Hausen H. 2009. Papillomaviruses in the causation of human cancers - a brief historical account. Virology 384:260–265.

2. Marur S, D’Souza G, Westra WH, Forastiere AA. 2010. HPV-associated head and neck cancer: a virus-related cancer epidemic. The LancetOncology 11:781–789.

3. Gillison ML, Koch WM, Capone RB, Spafford M, Westra WH, Wu L, Zahurak ML, Daniel RW, Viglione M, Symer DE, Shah KV, Sidransky D. 2000. Evidence for a causal association between human papillomavirus and a subset of head and neck cancers. Journal of the National Cancer Institute 92:709–720.

4. Ang KK, Harris J, Wheeler R, Weber R, Rosenthal DI, Nguyen-TÃ¢n PF, Westra WH, Chung CH, Jordan RC, Lu C, Kim H, Axelrod R, Silverman CC, Redmond KP, Gillison ML. 2010. Human papillomavirus and survival of patients with oropharyngeal cancer. The New England journal of medicine 363:24–35.

5. Gillison ML. 2004. Human papillomavirus-associated head and neck cancer is a distinct epidemiologic, clinical, and molecular entity. Seminars in oncology 31:744–754.

6. Songock WK, Kim SM, Bodily JM. 2017. The human papillomavirus E7 oncoprotein as a regulator of transcription. Virus research 231:56–75.

7. Pang CL, Toh SY, He P, Teissier S, Ben Khalifa Y, Xue Y, Thierry F. 2014. A functional interaction of E7 with B-Myb-MuvB complex promotes acute cooperative transcriptional activation of both S- and M-phase genes. (129 c). Oncogene 33:4039–4049.

8. Hoppe-Seyler K, Bossler F, Braun JA, Herrmann AL, Hoppe-Seyler F. 2018. The HPV E6/E7 Oncogenes: Key Factors for Viral Carcinogenesis and Therapeutic Targets. Trends in microbiology 26:158–168.

9. Mittal S, Banks L. 2017. Molecular mechanisms underlying human papillomavirus E6 and E7 oncoprotein-induced cell transformation. Mutation researchReviews in mutation research 772:23–35.

10. Howie HL, Koop JI, Weese J, Robinson K, Wipf G, Kim L, Galloway DA. 2011. Beta-HPV 5 and 8 E6 promote p300 degradation by blocking AKT/p300 association. PLoS pathogens 7:e1002211.

11. Muench P, Probst S, Schuetz J, Leiprecht N, Busch M, Wesselborg S, Stubenrauch F, Iftner T. 2010. Cutaneous papillomavirus E6 proteins must interact with p300 and block p53-mediated apoptosis for cellular immortalization and tumorigenesis. Cancer research 70:6913–6924.

12. Patel D, Huang SM, Baglia LA, McCance DJ. 1999. The E6 protein of human papillomavirus type 16 binds to and inhibits co-activation by CBP and p300. The EMBO journal 18:5061–5072.

13. Thomas MC, Chiang CM. 2005. E6 oncoprotein represses p53-dependent gene activation via inhibition of protein acetylation independently of inducing p53 degradation. Molecular cell 17:251–264.

14. White EA, Kramer RE, Tan MJ, Hayes SD, Harper JW, Howley PM. 2012. Comprehensive analysis of host cellular interactions with human papillomavirus E6 proteins identifies new E6 binding partners and reflects viral diversity. Journal of virology 86:13174–13186.

15. Zimmermann H, Degenkolbe R, Bernard HU, O’Connor MJ. 1999. The human papillomavirus type 16 E6 oncoprotein can down-regulate p53 activity by targeting the transcriptional coactivator CBP/p300. Journal of virology 73:6209–6219.

16. Xie X, Piao L, Bullock BN, Smith A, Su T, Zhang M, Teknos TN, Arora PS, Pan Q. 2014. Targeting HPV16 E6-p300 interaction reactivates p53 and inhibits the tumorigenicity of HPV-positive head and neck squamous cell carcinoma. Oncogene 33:1037–1046.

17. McBride AA. 2013. The Papillomavirus E2 proteins. Virology 445:57–79.

18. Bouvard V, Storey A, Pim D, Banks L. 1994. Characterization of the human papillomavirus E2 protein: evidence of trans-activation and trans-repression in cervical keratinocytes. The EMBO journal 13:5451–5459.

19. McBride AA, McPhillips MG, Oliveira JG. 2004. Brd4: tethering, segregation and beyond. Trends in microbiology 12:527–529.

20. Gauson EJ, Windle B, Donaldson MM, Caffarel MM, Dornan ES, Coleman N, Herzyk P, Henderson SC, Wang X, Morgan IM. 2014. Regulation of human genome expression and RNA splicing by human papillomavirus 16 E2 protein. Virology 468-470:10–18.

21. Sunthamala N, Thierry F, Teissier S, Pientong C, Kongyingyoes B, Tangsiriwatthana T, Sangkomkamhang U, Ekalaksananan T. 2014. E2 proteins of high risk human papillomaviruses down-modulate STING and IFN-kappa transcription in keratinocytes. PloS one 9:e91473.

22. Sunthamala N, Pang CL, Thierry F, Teissier S, Pientong C, Ekalaksananan T. 2014. Genome-wide analysis of high risk human papillomavirus E2 proteins in human primary keratinocytes. Genomics data 2:147–149.

23. Thierry Fo, Benotmane MA, Demeret C, Carcinogenesis P, Desaintes C. 2004. A Genomic Approach Reveals a Novel Mitotic Pathway in Papillomavirus Carcinogenesis A Genomic Approach Reveals a Novel Mitotic Pathway in.895–903.

24. Hadaschik D, Hinterkeuser K, Oldak M, Pfister HJ, Smola-Hess S. 2003. The Papillomavirus E2 protein binds to and synergizes with C/EBP factors involved in keratinocyte differentiation. Journal of virology 77:5253–5265.

25. Wu MH, Chan JY, Liu PY, Liu ST, Huang SM. 2007. Human papillomavirus E2 protein associates with nuclear receptors to stimulate nuclear receptor- and E2-dependent transcriptional activations in human cervical carcinoma cells. The international journal of biochemistry & cell biology 39:413–425.

26. Oldak M, Maksym RB, Sperling T, Yaniv M, Smola H, Pfister HJ, Malejczyk J, Smola S. 2010. Human papillomavirus type 8 E2 protein unravels JunB/Fra-1 as an activator of the beta4-integrin gene in human keratinocytes. Journal of virology 84:1376–1386.

27. Delcuratolo M, Fertey J, Schneider M, Schuetz J, Leiprecht N, Hudjetz B, Brodbeck S, Corall S, Dreer M, Schwab RM, Grimm M, Wu SY, Stubenrauch F, Chiang CM, Iftner T. 2016. Papillomavirus-Associated Tumor Formation Critically Depends on c-Fos Expression Induced by Viral Protein E2 and Bromodomain Protein Brd4. PLoS pathogens 12:e1005366.

28. Behren A, Simon C, Schwab RM, Loetzsch E, Brodbeck S, Huber E, Stubenrauch F, Zenner HP, Iftner T. 2005. Papillomavirus E2 protein induces expression of the matrix metalloproteinase-9 via the extracellular signal-regulated kinase/activator protein-1 signaling pathway. Cancer research 65:11613–11621.

29. Muhlen S, Behren A, Iftner T, Plinkert PK, Simon C. 2009. AP-1 and ERK1 but not p38 nor JNK is required for CRPV early protein 2-dependent MMP-9 promoter activation in rabbit epithelial cells. Virus research 139:100–105.

30. Muhlen S, Behren A, Iftner T, Simon C. 2010. Influence of HPV16 E2 and its localisation on the expression of matrix metalloproteinase-9. International journal of oncology 37:337–345.

31. Evans MR, James CD, Loughran O, Nulton TJ, Wang X, Bristol ML, Windle B, Morgan IM. 2017. An oral keratinocyte life cycle model identifies novel host genome regulation by human papillomavirus 16 relevant to HPV positive head and neck cancer. Oncotarget doi:10.18632/oncotarget.18328 [doi].

32. Cheon H, Holvey-Bates EG, Schoggins JW, Forster S, Hertzog P, Imanaka N, Rice CM, Jackson MW, Junk DJ, Stark GR. 2013. IFNbeta-dependent increases in STAT1, STAT2, and IRF9 mediate resistance to viruses and DNA damage. The EMBO journal 32:2751–2763.

33. Cheon H, Stark GR. 2009. Unphosphorylated STAT1 prolongs the expression of interferon-induced immune regulatory genes. Proceedings of the National Academy of Sciences of the United States of America 106:9373–9378.

34. Fink K, Grandvaux N. 2013. STAT2 and IRF9: Beyond ISGF3. Jak-Stat 2:e27521.

35. Chang YE, Laimins LA. 2000. Microarray analysis identifies interferon-inducible genes and Stat-1 as major transcriptional targets of human papillomavirus type 31. Journal of virology 74:4174–82.

36. Rincon-Orozco B, Halec G, Rosenberger S, Muschik D, Nindl I, Bachmann A, Ritter TM, Dondog B, Ly R, Bosch FX, Zawatzky R, RÃ¶sl F. 2009. Epigenetic silencing of interferon-kappa in human papillomavirus type 16-positive cells. Cancer research 69:8718–25.

37. Kaczkowski B, Morevati M, Rossing M, Cilius F, Norrild B. 2012. A Decade of Global mRNA and miRNA Profiling of HPV-Positive Cell Lines and Clinical Specimens. The open virology journal 6:216–31.

38. Kaczkowski B, Rossing M, Andersen DK, Dreher A, Morevati M, Visser Ma, Winther O, Nielsen FC, Norrild B. 2012. Integrative analyses reveal novel strategies in HPV11,-16 and −45 early infection. Scientific reports 2:515–515.

39. Reiser J, Hurst J, Voges M, Krauss P, Munch P, Iftner T, Stubenrauch F. 2011. High-risk human papillomaviruses repress constitutive kappa interferon transcription via E6 to prevent pathogen recognition receptor and antiviral-gene expression. Journal of virology 85:11372–11380.

40. Li S, Labrecque S, Gauzzi MC, Cuddihy AR, Wong AH, Pellegrini S, Matlashewski GJ, Koromilas AE. 1999. The human papilloma virus (HPV)-18 E6 oncoprotein physically associates with Tyk2 and impairs Jak-STAT activation by interferon-alpha. Oncogene 18:5727–5737.

41. Barnard P, McMillan NA. 1999. The human papillomavirus E7 oncoprotein abrogates signaling mediated by interferon-alpha. Virology 259:305–313.

42. Barnard P, Payne E, McMillan NA. 2000. The human papillomavirus E7 protein is able to inhibit the antiviral and anti-growth functions of interferon-alpha. Virology 277:411–419.

43. Ramqvist T, Mints M, Tertipis N, Nasman A, Romanitan M, Dalianis T. 2015. Studies on human papillomavirus (HPV) 16 E2, E5 and E7 mRNA in HPV-positive tonsillar and base of tongue cancer in relation to clinical outcome and immunological parameters. Oral oncology 51:1126–1131.

44. Anayannis NV, Schlecht NF, Ben-Dayan M, Smith RV, Belbin TJ, Ow TJ, Blakaj DM, Burk RD, Leonard SM, Woodman CB, Parish JL, Prystowsky MB. 2018. Association of an intact E2 gene with higher HPV viral load, higher viral oncogene expression, and improved clinical outcome in HPV16 positive head and neck squamous cell carcinoma. PloS one 13:e0191581.

45. Nulton TJ, Kim NK, DiNardo LJ, Morgan IM, Windle B. 2018. Patients with integrated HPV16 in head and neck cancer show poor survival. Oral oncology 80:52–55.

46. Nulton TJ, Olex AL, Dozmorov M, Morgan IM, Windle B. 2017. Analysis of the cancer genome atlas sequencing data reveals novel properties of the human papillomavirus 16 genome in head and neck squamous cell carcinoma. Oncotarget 8:17684–17699.

47. Smith JA, White EA, Sowa ME, Powell ML, Ottinger M, Harper JW, Howley PM. 2010. Genome-wide siRNA screen identifies SMCX, EP400, and Brd4 as E2-dependent regulators of human papillomavirus oncogene expression. Proceedings of the National Academy of Sciences of the United States of America 107:3752–3757.

48. Kuss-Duerkop SK, Westrich JA, Pyeon D. 2018. DNA Tumor Virus Regulation of Host DNA Methylation and Its Implications for Immune Evasion and Oncogenesis. Viruses 10:10.3390/v10020082.

49. Momparler RL, Cote S, Eliopoulos N. 1997. Pharmacological approach for optimization of the dose schedule of 5-Aza-2’-deoxycytidine (Decitabine) for the therapy of leukemia. Leukemia 11:175–180.

50. Dickson MA, Hahn WC, Ino Y, Wu JY, Weinberg RA, Louis DN, Li FP, Rheinwald JG, Ronfard V. 2000. Human Keratinocytes That Express hTERT and Also Bypass a p16INK4a-Enforced Mechanism That Limits Life Span Become Immortal yet Retain Normal Growth and Differentiation Characteristics Human Keratinocytes That Express hTERT and Also Bypass a p16 INK4a-Enf. doi:10.1128/MCB.20.4.1436-1447.2000.Updated.

51. Xue Y, Bellanger S, Zhang W, Lim D, Low J, Lunny D, Thierry F. 2010. HPV16 E2 is an immediate early marker of viral infection, preceding E7 expression in precursor structures of cervical carcinoma. Cancer research 70:5316–5325.

52. Vosa L, Sudakov A, Remm M, Ustav M, Kurg R. 2012. Identification and analysis of papillomavirus E2 protein binding sites in the human genome. Journal of virology 86:348–357.

53. Jang MK, Shen K, McBride AA. 2014. Papillomavirus genomes associate with BRD4 to replicate at fragile sites in the host genome. PLoS pathogens 10:e1004117.

54. Gauson EJ, Wang X, Dornan ES, Herzyk P, Bristol M, Morgan IM. 2016. Failure to interact with Brd4 alters the ability of HPV16 E2 to regulate host genome expression and cellular movement. Virus research 211:1–8.

55. Terenzi F, Saikia P, Sen GC. 2008. Interferon-inducible protein, P56, inhibits HPV DNA replication by binding to the viral protein E1. The EMBO journal 27:3311–3321.

56. Saikia P, Fensterl V, Sen GC. 2010. The inhibitory action of P56 on select functions of E1 mediates interferon’s effect on human papillomavirus DNA replication. Journal of virology 84:13036–13039.

57. Khodarev NN, Beckett M, Labay E, Darga T, Roizman B, Weichselbaum RR. 2004. STAT1 is overexpressed in tumors selected for radioresistance and confers protection from radiation in transduced sensitive cells. Proceedings of the National Academy of Sciences of the United States of America 101:1714–1719.

58. Weichselbaum RR, Ishwaran H, Yoon T, Nuyten DS, Baker SW, Khodarev N, Su AW, Shaikh AY, Roach P, Kreike B, Roizman B, Bergh J, Pawitan Y, van de Vijver MJ, Minn AJ. 2008. An interferon-related gene signature for DNA damage resistance is a predictive marker for chemotherapy and radiation for breast cancer. Proceedings of the National Academy of Sciences of the United States of America 105:18490–18495.

59. Piboonniyom SO, Duensing S, Swilling NW, Hasskarl J, Hinds PW, Munger K. 2003. Abrogation of the retinoblastoma tumor suppressor checkpoint during keratinocyte immortalization is not sufficient for induction of centrosome-mediated genomic instability. Cancer research 63:476–483.

60. Bentley P, Tan MJA, McBride AA, White EA, Howley PM. 2018. The SMC5/6 complex interacts with the papillomavirus E2 protein and influences maintenance of viral episomal DNA. J Virol doi:10.1128/jvi.00356-18.

61. White EA, Sowa ME, Tan MJ, Jeudy S, Hayes SD, Santha S, Munger K, Harper JW, Howley PM. 2012. Systematic identification of interactions between host cell proteins and E7 oncoproteins from diverse human papillomaviruses. Proceedings of the National Academy of Sciences of the United States of America 109:E260–7.

